# *LADON*, a natural antisense transcript of *NODAL*, promotes an invasive behaviour in melanoma cells

**DOI:** 10.1101/2020.04.09.032375

**Authors:** Dutriaux Annie, Diazzi Serena, Caburet Sandrine, Bresesti Chiara, Hardouin Sylvie, Deshayes Frédérique, Collignon Jérôme, Flagiello Domenico

## Abstract

The TGFβ family member NODAL, known for its role during embryonic development, has also been associated with tumor progression in several cancers. Some of the evidence supporting its involvement in melanoma appeared contradictory, suggesting that NODAL in this context might rely on a non-canonical signalling mode. We found that NODAL inactivation in a metastatic melanoma cell line prevents the cells from acquiring invasive behaviour. However, we show that this phenotype does not result from the absence of NODAL, but from a defect in the expression of a natural antisense transcript of NODAL, here called LADON. We found that LADON promotes the mesenchymal to amoeboid transition that is critical to melanoma cell invasiveness, and that a WNT/β-CATENIN signalling-dependent increase in LADON expression is required to complete this transition. LADON’s downstream effectors include, among others, the proto-oncogene MYCN. These results identify LADON as a player in the regulatory network that governs tumor progression in melanoma, and possibly in other types of cancer.

## Introduction

Metastasis is responsible for up to 90% of cancer deaths (Chaffer & Weinberg, 2011), making the determination of the molecules involved in this process crucial to the improvement of diagnosis and treatment. The *NODAL* gene, which encodes a TGF*β* family member, was identified as a possible candidate (Quail et al., 2013; Topczewska et al., 2006). The NODAL ligand binds a receptor complex containing type I (ALK4 or 7) and type II (ActRIIA or B) serine/threonine kinase receptors, the activation of which leads to the phosphorylation of the signal transducers SMAD2 or SMAD3 (Massagué et al., 2005). NODAL is best known for its role during development, where it is required both to maintain the undifferentiated state of embryonic precursors and to specify the identity of specific cell types, including several motile cell types (Robertson, 2014). It is also expressed in the adult, notably in tissues that undergo periodic renewal or remodelling under the control of hormonal stimuli, such as the endometrium and the mammary gland (Bianco et al., 2002; Papageorgiou et al., 2009). NODAL expression in tumor cells has been correlated with their plasticity and invasive behaviour (Bodenstine et al., 2016; Quail et al., 2013), in line with its functions in embryonic and adult tissues.

The involvement of NODAL in tumor progression and metastasis was first described in melanoma cell lines (Topczewska et al., 2006), but although evidence supporting such a role for this factor has been obtained in other cancer types, its actual contribution to metastasis in melanoma has been disputed (Donovan et al., 2017; Hooijkaas et al., 2011), as its expression was not always detected there. A review of available *NODAL* expression data in cancer cells suggested the existence of distinct splice variants (Strizzi et al., 2012), all including the second exon of the gene but sometimes, notably in melanoma cell lines, lacking its third (and last) exon and therefore the capacity to produce a functional ligand. A previous study had already shown that the expression of CRIPTO, an obligatory co-receptor of NODAL, was weak in metastatic melanoma cell lines, and restricted to a small cell subpopulation (Postovit et al., 2008). These observations raised the possibility in this context of a non-canonical mode of action of *NODAL*, not necessarily reliant on its known signalling function.

We thus used both genetic and pharmacological approaches to reassess the importance of *NODAL* in mediating the invasive properties of melanoma cell lines. Our results argue that the NODAL protein plays no role, but that a natural antisense transcript, which we have named LADON, which overlaps with *NODAL* exon2, responds to signals controlling tumor progression and metastasis and regulates the expression of known oncogenes and tumor suppressors. Our study identifies *LADON* as a novel regulator of the metastatic process.

## Results

### The invasive behaviour of A375 cells is dependent on the presence of *NODAL* exon2

We used RT-PCR analysis to characterize *NODAL* expression in a panel of human cell lines relevant to cancer and melanoma: melanocytes, non-metastatic melanoma (MNT1), metastatic melanoma (A375, FO1, 888Mel, SLM8), breast cancer (MCF7), embryonic kidney (HEK293). A primer pair specific for *NODAL* exon2 amplified the expected band in all of them (Fig 1A, S1A). In contrast, a primer pair spanning the junction of exon2 and 3 (Fig 1A) only detected a transcript in the embryonic kidney cell line (Fig S1B). These results are consistent with the presence of exon2-containing transcripts and the absence of full-length *NODAL* transcripts, as has been reported in other melanoma cell lines (Donovan et al., 2017; Strizzi et al., 2012)

**Figure 1.**
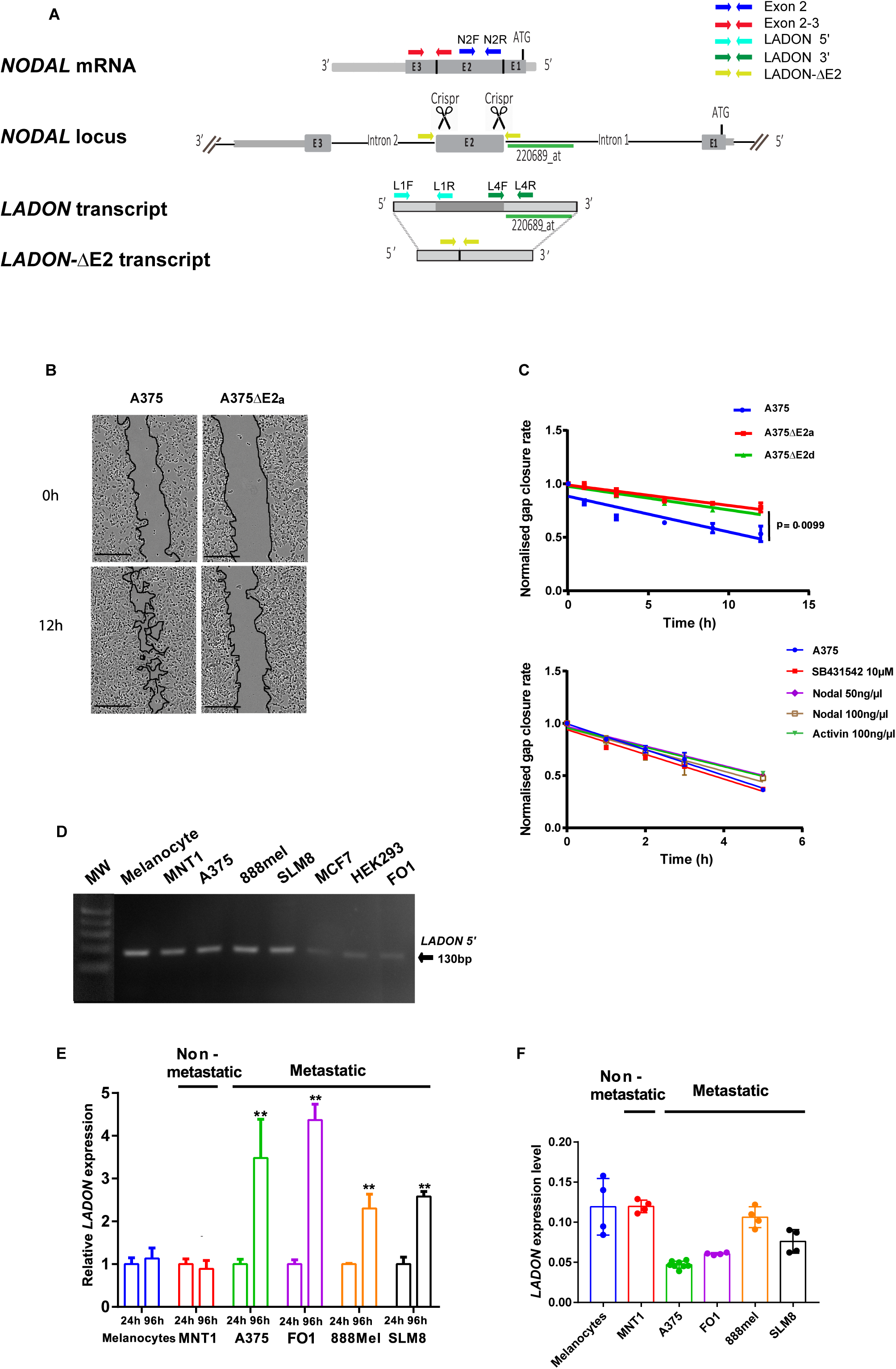
The invasive behavior of A375 cells is dependent on *LADON*, not on NODAL. A Schematic of the human *NODAL* locus with its 3 exons (E1 to E3), showing the full-length *NODAL* mRNA above, and the *LADON* transcript transcribed from the opposite strand below. The truncated *LADON*-ΔE2 transcript is expressed in A375 cells deleted for *NODAL* exon2. The primers used to track these transcripts are represented by arrows. B Representative images, acquired at t=0 and t=12h, of scratch-wound healing assays performed with A375 and A375ΔE2a cells. C Top: Normalized gap closure rates of A375, A375ΔE2a and A375ΔE2d cells during the scratch-wound healing assay shown in (B). Bottom: Normalized gap closure rates of A375 cells treated with or without recombinant NODAL, recombinant ACTIVIN or the ACTIVIN/NODAL pathway inhibitor SB431542. The p-value was calculated by a two-way ANOVA test. D RT-PCR with primers (L1F-L1R) amplifying a 130nt band from the 5’ region of *LADON* detects the presence of the transcript in all cell lines of the panel: melanocytes, non-metastatic melanoma (MNT1), metastatic melanoma (A375, 888mel, SLM8, FO1), breast cancer (MCF7), embryonic kidney (HEK293). E RT-qPCR measurements of *LADON* expression were performed in melanocytes, non-metastatic (MNT1) or metastatic melanoma cells (A375, 888Mel, SLM8 and FO1), after 24h and 96h of culture. *LADON* expression was normalized to that of endogenous *RPL13*. For each cell line the value at 24h was then set to 1. F RT-qPCR measurements of *LADON* expression after 24h culture performed on the same panel as in (E), but just normalized to endogenous *RPL13* expression. Histograms display mean values ± SD from a minimum of two independent replicates. P-values were calculated by Student’s t test, ** < 0.01.

To test for a possible *NODAL* exon2 requirement during key steps of the metastatic process we used genome editing to delete it in the three copies of the gene present in A375 cells (Fig 1A). Five of the independent mutant clones thus obtained, designated A375*Δ*E2a to e, were characterized (Fig S1C, D). The absence of *NODAL* exon2 was found to impair their ability to close the gap in a 2D wound-healing (scratch) assay (Fig 1B, C). This effect was detected in less than 12h of culture, which suggests that it results from a decrease in cell motility. However, increasing concentrations of recombinant NODAL or ACTIVIN, another TGF*β*-related ligand that signals via the same receptor complex as NODAL, had no effect on the behavior of A375 cells in the same assay (Fig 1C). Treatment with SB431542, a pharmacological inhibitor of the ACTIVIN/NODAL type I receptors ALK4, 5 and 7, also had no effect (Fig 1C).

These results imply that A375 cells require *NODAL* exon2 to adopt their invasive phenotype, but their lack of response to signalling molecules and inhibition of the ACTIVIN/NODAL signalling pathway, as well as the absence of full-length *NODAL* transcripts, suggest that the mode of action of the gene does not involve the usual signalling function of its gene product.

### A natural antisense transcript overlaps with *NODAL* exon2 in melanoma cell lines

Consultation of the *Ensembl* database (GRCh38.p13 primary assembly) revealed the presence of a third transcript at the human *NODAL* locus in addition to the two *NODAL* splice variants already known (Fig 1A, S1E). This 1728nt long RNA (AC022532) is transcribed from the plus strand (opposite to *NODAL*). The corresponding cDNA was characterized as being full-length (Ota et al., 2004). It starts in *NODAL* intron2 and ends in *NODAL* intron1, and therefore includes a sequence complementary to that of the entire *NODAL* exon2. This natural antisense transcript (NAT) contains an ORF, but lacks a proper Kozak consensus sequence at the ATG and no corresponding protein has been reported so far. It is annotated as non-coding. Homologous transcripts of similar size were found in Chimpanzee and in Orangutan (Ensembl database). A longer homolog has been found in pigs, but none has been reported in mice and rats, nor in any of the other classical vertebrate animal models (zebrafish, xenopus, chick), which may suggest that it arose within the mammalian lineage.

To confirm the presence of this antisense transcript in A375 cells we reverse-transcribed total A375 RNA using either a reverse (N2R) or a forward (N2F) primer with respect to the expected orientation of its transcription. The resulting cDNAs were used to PCR-amplify sequences spanning exon2 and the adjoining 5’ or 3’ regions (Fig 1A). The fact that a band of the expected size was only obtained with cDNAs transcribed using the reverse primer N2R confirmed the absence of *NODAL* expression and demonstrated the presence of the antisense transcript (Fig S1F). The corresponding transcription unit is designated as *LADON* hereafter.

To get a sense of transcription patterns at the *NODAL* locus we compared strand specific RNA-seq data (ENCODE project) at this position for a number of cell lines (Fig S1E). Plus strand transcription is far more prevalent than minus strand transcription, as instances of misidentified *NODAL* expression led us to expect. Both strands are expressed in human ESCs and in keratinocytes for example, but only plus strand expression is detected in melanocytes and in melanoma cell lines such as the A375 line. While reads originating from minus strand transcription are neatly positioned over *NODAL* exons, reads originating from the plus strand cover an area that includes *LADON* but is much broader, suggesting that below a certain level *LADON* expression cannot be distinguished from surrounding pervasive expression.

We thus used a specific pair of primers (L1F-L1R, Fig 1A) to detect *LADON* transcripts, and amplified a band of the expected size in all the cell lines of our panel (Fig 1D). RT-qPCR analysis of *LADON* expression in melanocyte and melanoma cell lines showed big variations in the expression of *LADON* within individual metastatic melanoma cell lines but not in melanocyte and non-metastatic cell lines. The extent of these variations was correlated to the length of time the cells had been cultured, as a 2- to 4-fold increase in *LADON* expression was observed between 24h and 96h of culture (Fig 1E). Interestingly, basal *LADON* expression was lower in metastatic melanoma cells than in melanocytes and non-metastatic melanoma cells (Fig 1F), and the lower this expression, the higher the subsequent increase in *LADON* expression after 96h (Fig 1E). In contrast, A375*Δ*E2 clones, which expressed a truncated *LADON* transcript (*LADON-ΔE2*) reduced to the 5’ and 3’ parts that flank exon2 (Fig 1A, S1G), showed no sign of increased expression after 96h in culture, implying that the deletion deprived *LADON* of a critical regulatory input (Fig S1H). The capacity of melanoma cell lines to metastasize therefore appears to correlate with their capacity to increase *LADON* expression.

These results confirmed that the absence of *NODAL* did not cause the changes in behaviour we characterized in cells deleted for *NODAL* exon2. They led us to identify *LADON* as a candidate for this cell phenotype, and to investigate the link between the common propensity of metastatic cell lines to up-regulate *LADON* and to adopt invasive behaviour.

### Melanoma cells secrete factors that promote *LADON* expression

To investigate what triggered the increase in *LADON* expression in the A375 cell line, we first tested its dependence on cell density (Kim et al., 2017). *LADON* transcript levels measured after 24h culture in cells seeded at high (80%) density were actually no different from those measured in cells seeded at low (20%) density (Fig 2A). The increase in *LADON* expression found after 72h culture was therefore more likely to depend on the duration of the culture. To assess the possible dependency of *LADON* expression on secreted factors accumulating in the culture medium A375 cells were grown in standard medium (SM), with or without 10% fetal calf serum (FCS). The media were collected three days later when the cells reached high density. These A375-conditioned media (CM) were then used to culture cells seeded at low density for 24h. For A375 cells these treatments resulted in a 2.4-fold increase in the expression of *LADON* (Fig 2B), compared to its expression level when cultured in SM. This increase occurred regardless of the presence of FCS, thus eliminating nutrient depletion as a possible cause (Fig S2A). Another metastatic melanoma cell line, FO1, was also found to increase *LADON* expression when cultured in its own CM (Fig S2B). In contrast, cells from the non-tumorigenic non-metastatic MNT1 cell line did not increase *LADON* expression when exposed to their own CM or to that of A375 cells (Fig 2B), but this MNT1 CM did trigger an increase in *LADON* expression in A375 cells. So, while both A375 and MNT1 cells secrete and accumulate *LADON*-inducing factors in their culture medium, only A375 cells are able to respond accordingly, suggesting that the ability to respond to *LADON*-inducing factors is specific to metastatic cell lines.

**Figure 2.**
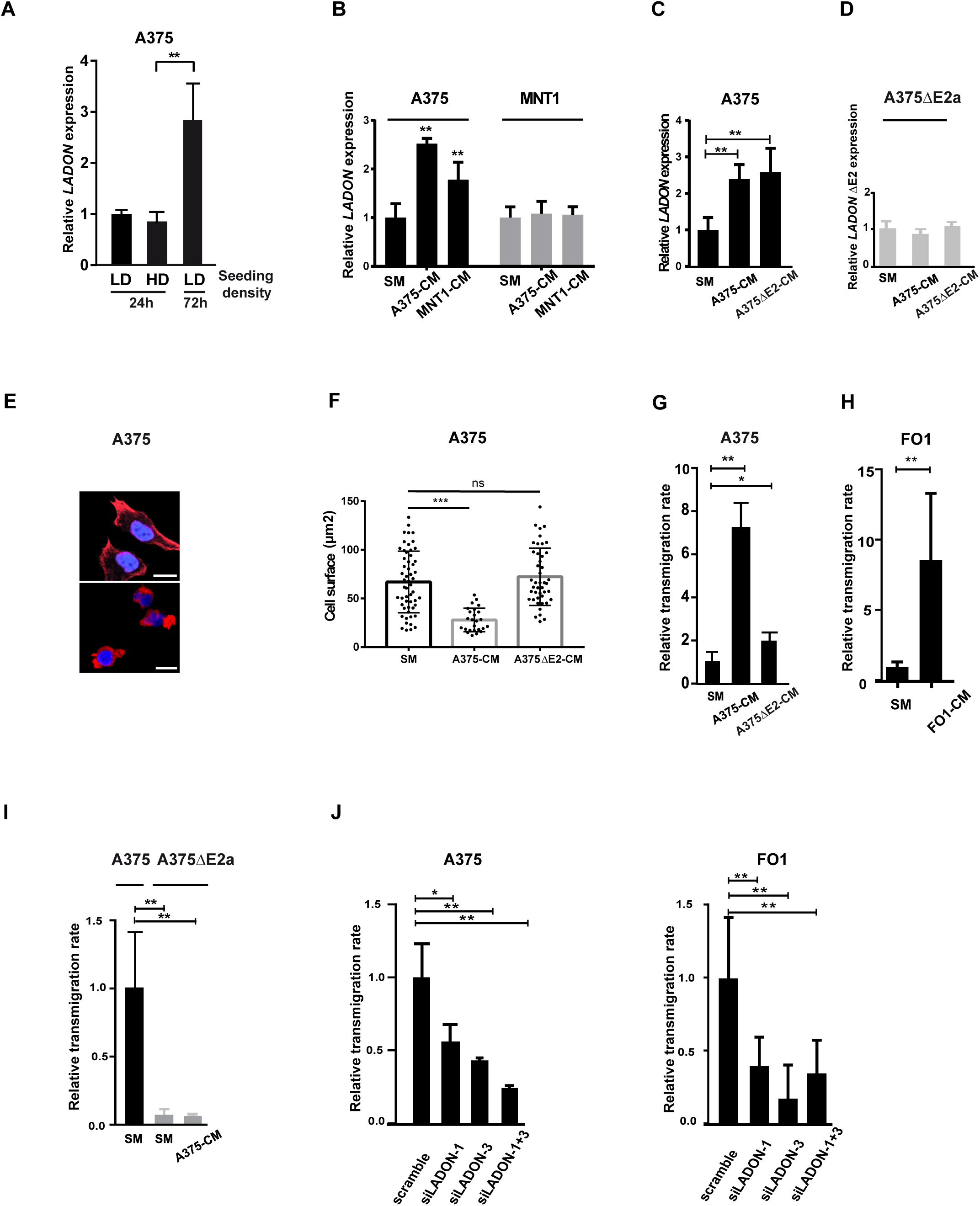
Melanoma cells secrete factors that promote *LADON* expression, and invasion. A RT-qPCR analysis of *LADON* expression in A375 cells seeded at low (LD) or high density (HD) and cultured for 24 or 72h. Cell density does not affect LADON’s expression level whereas cell culture duration does. Values were normalized to that of the LD condition after 24h, which was set to 1. B RT-qPCR analysis of *LADON* expression in A375 and MNT1 cells cultured for 24h in standard culture medium (SM), A375-conditioned medium (A375-CM) or MNT1- conditioned medium (MNT1-CM). Values were normalized to that of the SM condition, which was set at 1. C,D RT-qPCR analysis of *LADON* and *LADON-ΔE2* expression in A375 or A375ΔE2a cells cultured for 24h in SM, A375-CM or A375ΔE2-CM. Values were normalized to that of the SM condition, which was set to 1. E Representative images of A375 cells with elongated or rounded morphology, visualized after F-actin (red) and Hoechst 33342 (nuclear, blue) staining. Scale bar 25µm. F Histogram of cell surface measurements of A375 cells cultured for 24h in SM, A375-CM or A375ΔE2-CM (n>20) from at least 3 independent experiments. G, H, I Relative transmigration rates of A375, FO1 or A375ΔE2a cells cultured in SM, A375-CM, FO1-CM or A375ΔE2-CM. Values were normalized to that of the SM condition, which was set to 1. I Relative transmigration rates of A375 or FO1 cells in the presence of the siRNAs scrambled (a control), siLADON-1, siLADON-3, or siLADON-1 and -3. Values were normalized to that obtained with the scrambled siRNA, which was set to 1. Histograms display mean values ± SD from a minimum of three independent replicates. P-values were calculated in a Student’s t test, * < 0.05, ** < 0.01.

Analysis of the published content of A375 cell-derived exosomes (GSE35386; (Xiao et al., 2012) revealed a relative enrichment in *LADON* transcripts (Fig S2C). This raised the question of their possible contribution to the inductive capacity of the CM. We therefore analyzed the effect on A375 cells of a CM obtained from A375*Δ*E2 cells (A375*Δ*E2-CM). This exon2-depleted CM had the same capacity to induce *LADON* expression as that obtained from A375 cells (Fig 2C), implying that a full-length *LADON* is not required in the CM to increase its own expression. Cells from A375*Δ*E2 clones, however, failed to increase the expression of the *LADON-ΔE2* transcript when exposed to the same CM (Fig 2D), confirming that the capacity of the locus to respond to these treatments is dependent on its integrity.

### *LADON* expression promotes invasion

Migrating melanoma cells rely on two inter-convertible modes of migration, designated mesenchymal and amoeboid (Friedl & Wolf, 2010; Sanz-Moreno et al., 2008) with the faster amoeboid mode being the most efficient to drive tumor metastasis (Gadea et al., 2007; Panková et al., 2010; Sanz-Moreno et al., 2011). We therefore investigated the effect of the A375-CM on cell morphology and migratory behaviour. The mesenchymal to amoeboid transition (MAT), which involves a change from an elongated to a smaller and rounded cell morphology, was monitored via F-actin staining (Fig 2E) (Sahai & Marshall, 2003). In SM no clear difference was seen between cultures of mutant and unmodified A375 cell populations, both showing similar cell size ranges (Fig S2D). However, when grown in CM, the average size of A375 cells decreased, with cells covering an area greater than 50 *μ*m^2^ disappearing, and their roundness index increased (Fig 2F and S2E). When the capacity of these cells to transmigrate through a collagen layer was quantified over a 24h period, this change in the composition of the population was associated with a 7-fold increase in the rate of transmigration (Fig 2G). FO1 cells grown in their own CM showed a similar decrease in mean cell size and an increase in their transmigration rate (Fig S2F and 2H). In contrast, A375*Δ*E2 cells, whose transmigration rate in SM was one-tenth that of A375 cells, showed no improvement in transmigration capacity when cultured in CM (Fig 2I). *LADON* is therefore endowing A375 cells with the capacity to undergo CM-induced changes in cell behaviour. Again, to assess the contribution of exon2 sequences to this particular effect of the CM we analyzed the effect of the A375*Δ*E2-CM on A375 cells. Although this treatment had no significant impact on the cell size of A375 cells (Fig 2F) their transmigration rate showed a slight increase (Fig 2G). The impact of this CM was therefore far less important than that from A375 cells, which therefore appears to be crucially dependent on the presence of exon2 in the CM-producing cells.

To confirm that it is the depletion of the *LADON* transcript and not another unrelated consequence of the deletion that is causing invasiveness to drop, we examined how knocking-down *LADON* expression affects the behaviour of A375 and FO1 cells. Three different siRNAs located at the 5’ end of the gene were tested. RT-PCR results showed a significant decrease of *LADON* expression with two of them, used alone or together (Fig S2G). The knockdown (KD) of *LADON* severely impaired the ability of A375 and FO1 cells to transmigrate through a collagen monolayer over 24h, a behaviour reminiscent of that of A375*Δ*E2 clones (Fig 2J). We also noticed that in both cell lines *LADON* KD cells systematically reached confluence earlier than controls. They indeed had higher cell counts at the end of the culture (Fig S2H), suggesting that their proliferation rate was positively impacted, an observation consistent with these cells maintaining a proliferative and non-invasive behavior(Carreira et al., 2006; Hoek et al., 2008).

Taken together, these results show that the expression of *LADON* in melanoma cells conditions both their capacity to produce factors that promote changes in cell behaviour conducive to metastasis (i.e., MAT, increase in motility, decrease in proliferation), and their capacity to respond to such factors.

### The increase in *LADON expression* promotes an increase in MYCN expression

To identify factors acting downstream of *LADON*, we used mass spectrometry to compare the proteomes of A375 and A375*Δ*E2 cells. The results are displayed in a volcano plot (Fig 3A). We focused on proteins that were identified by at least 2 independent peptides and with a fold change superior to 1.5. A Gene Ontology (GO) term analysis of the 28 proteins thus obtained, showed a significant enrichment in components of filamentous actin (PDLIM4, FREMT2 and NCKAP1), stress fibers (PDLIM4, Dynactin and FREMT2), as well as lamellipodia (PDLIM4, PLCG1, FREMT2 and NCKAP1) (Fig 3A; Table 1A). All of these were up-regulated in A375*Δ*E2 cells, suggesting a repressive role of *LADON* on their expression. Among the six proteins that had a fold change superior to 2 (Table 1B), 5 were overexpressed in A375*Δ*E2 cells and turned out to be known tumor (FAM107B, ACPP) or metastasis (NDRG1) suppressors (Guo et al., 2017; Meeusen & Janssens, 2018; Sharma et al., 2017), or transcription factors known to activate these proteins (p65, also known as NF-*κ*B regulatory subunit RELA, is an activator of ACPP) (Zelivianski et al., 2004). Conversely, the under-expressed protein PPP4R2 is a regulatory subunit of PPP4C, which is known to promote cancer cell growth and invasion (Hastie et al., 2000; X. Li et al., 2015). We focused our attention on NDRG1 (N-MYC downstream regulated gene 1), primarily known as a metastasis suppressor in various cancers (Sun, Zhang, Bae, et al., 2013).

**Figure 3.**
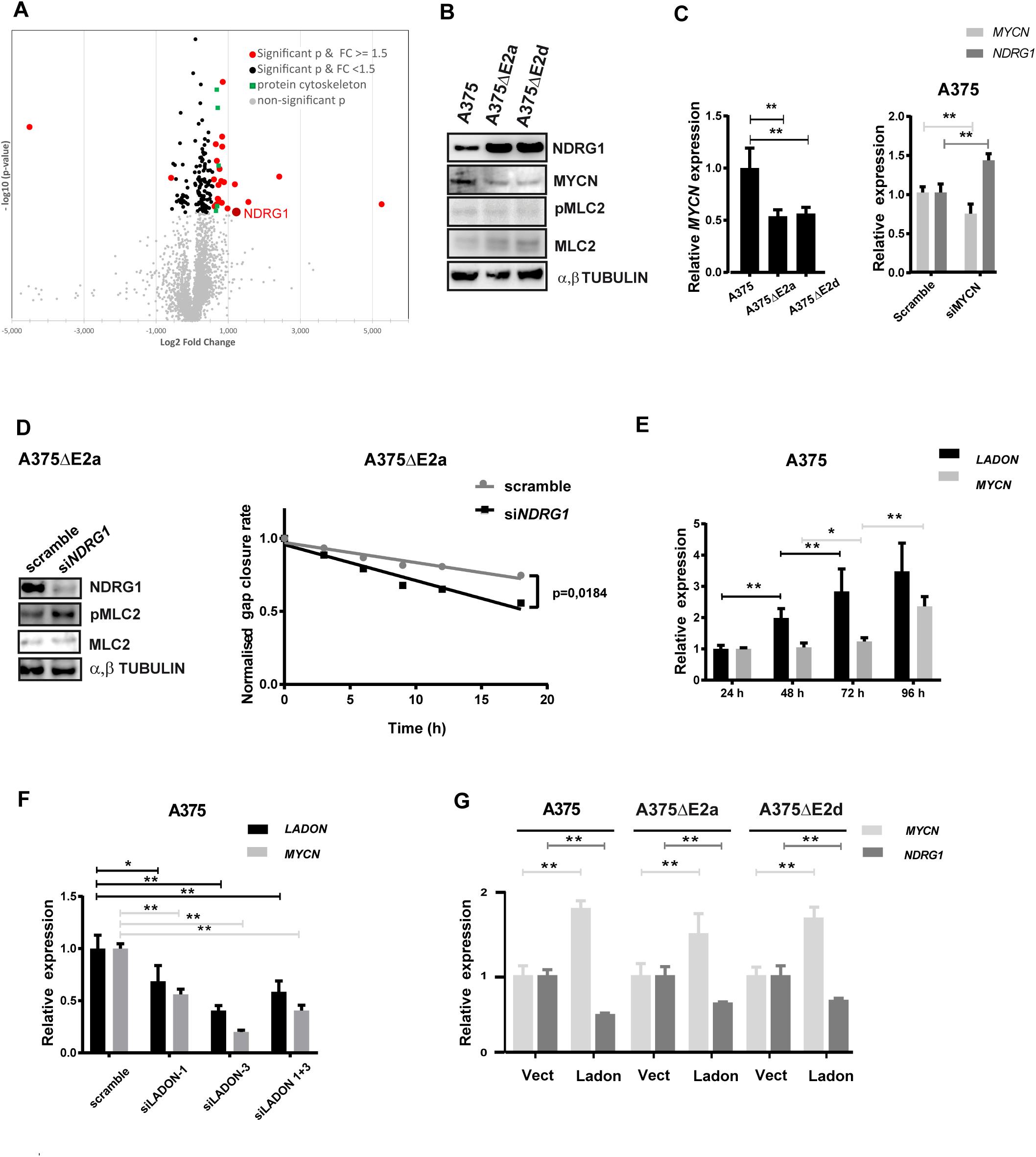
*LADON* promotes the expression of the proto-oncogene MYCN in A375 cells. A Volcano plot representation of a comparative proteomics analysis between A375 and A375ΔE2 cells. Grey dots: non-significant p-value. Black dots, significant p-value and fold change < 1,5. Red dots: significant p-value and fold change > 1,5. See also Table 1. B Western blot analysis of lysates from A375, A375ΔE2a and A375ΔE2d cells detecting NDRG1, MYCN, pMLC2, MCL2 and α,β-TUBULIN levels. C Left panel: RT-qPCR analysis of *MYCN* expression in A375 cells and A375ΔE2a cells. Right panel: RT-qPCR analysis of *MYCN* and *NDRG1* in A375 cells treated with scrambled or siMYCN siRNA. Expression levels were normalized to that of endogenous *RPL13*. Values were then normalized to those obtained for A375 cells or with the scrambled siRNA, which were set to 1. D Western blot analysis of lysates from A375ΔE2a cells treated with siRNAs scrambled or siNDRG1. NDRG1, pMLC2, MCL2 and α-β TUBULIN levels are revealed. Normalized gap closure rates of A375ΔE2a cells treated with the same siRNAs in a scratch-wound healing assays performed for18h. The p-values were calculated via a two-way ANOVA test. E RT-qPCR analysis of the expression dynamics of *LADON* and *MYCN* in A375 cells over a 96h culture. Values were normalized to those obtained at t=24h, which were set to 1. F RT-qPCR analysis of *LADON* and *MYCN* expression in A375 cells treated with siRNAs scrambled, siLADON-1, siLADON-3, or siLADON-1 and -3. Values were normalized to those obtained with the scrambled siRNA, which were set to 1. G RT-qPCR analysis of *MYCN* and *NDRG1* expression in A375, A375ΔE2a and A375ΔE2d cells transfected with *GFP* alone (vector) or together with *LADON*. Values were normalized to those obtained for cells treated with the vector alone, which were set to 1. Histograms display mean values ± SD from a minimum of three independent replicates. P-values were calculated by Student’s t test, * < 0.05.

**Table 1.** Differentially expressed proteins between A375 cells and *LADON* depleted A375ΔE2 cells. **(A)** Gene Ontology (GO) term analysis of differentially expressed proteins thus obtained showed a significant enrichment in 5 proteins involved in the composition of filamentous actin. **(B)** Proteins with a fold change superior to 2. Proteins were designated significant based on cutoff p value <0.05

| Max fold change | Description | Filamentous Actin | Stress Fiber | Lamellipodium | Highest mean condition | Accession | Anova (p) |
| --- | --- | --- | --- | --- | --- | --- | --- |
| 1,606767329 | PDZ and LIM domain protein 4 (PDLIM4) | X | X | X | Mutant | P50479 | 0,0016411 |
| 1,637021061 | 1-phosphatidylinositol 4,5-bisphosphate phosphodiesterase gamma-1 (PLCG1) |  |  | X | Mutant | P19174 | 0,00271601 |
| 1,664529632 | Dynactin subunit 4 (DCTN4) |  | X |  | Mutant | Q9UJW0 | 0,01351308 |
| 1,618822307 | Fermitin family homolog 2 (FERMT2) | X | X | X | Mutant | Q96AC1 | 0,0415036 |
| 1,580463985 | Nck-associated protein 1 (NCKAP1) | X |  | X | Mutant | Q9Y2A7 | 0,047219 |

| Max fold change | Description | Fonction | Highest mean condition | Accession | Anova (p) |
| --- | --- | --- | --- | --- | --- |
| 2,274639991 | Protein FAM107B (FAM107B) | tumor suppressor | Mutant | Q9H098 | 0,02268322 |
| 38,17912767 | Prostatic acid phosphatase (ACPP) | tumor suppressor, regulated by p65 | Mutant | P15309 | 0,03938661 |
| 22,78074964 | Serine/threonine-protein phosphatase 4 regulatory subunit 2 (PPP4R2) | overexpression in cancer promotes cell growth and invasion | WT | Q9NY27 | 0,00461911 |
| 5,3358974 | Transcription factor p65 (RELA) | transcription factor, cell proliferation, apoptosis and oncogenesis | Mutant | Q04206 | 0,01828842 |
| 2,33790023 | Protein NDRG1 (NDRG1) | metastasis suppressor | Mutant | Q92597 | 0,04885314 |
| 2,947432444 | Gamma-aminobutyric acid receptor-associated protein-like 2 (GABARAPL2) | Ubiquitin-like modifier involved in intra-Golgi traffic | Mutant | P60520 | 0,03702373 |

Western blot (WB) analyses confirmed that A375*Δ*E2 cells expressed higher levels of NDRG1 than A375 cells (Fig 3B). This increase correlated with lower levels of MYCN and lower levels of phosphorylated myosin light chain 2 (pMLC2). This was in line with studies showing that in prostate and colorectal cancer cells NDRG1 is negatively regulated by MYCN (J. Li & Kretzner, 2003) and inhibits actin filament polymerization, stress fibre assembly and cell migration via the ROCK1/pMLC2 pathway (Sun, Zhang, Zheng, et al., 2013). RT-PCR analyses found a significant reduction in *MYCN* expression in A375*Δ*E2 clones, and the KD of *MYCN* expression in unmodified A375 cells resulted in a significant increase in *NDRG1* expression (Fig 3C), as in A375*Δ*E2 cells. In contrast, the KD of *NDRG1* in A375*Δ*E2 cells resulted in higher pMLC2 levels and a rescue of their motility (Fig 3D). Much of the impact of *LADON* on cell behaviour therefore appears to be mediated by *MYCN*, a well-known proto-oncogene (Ruiz-Pérez et al., 2017), via its effect on MYCN downstream targets such as NDRG1.

To assess the implication of the *LADON* transcript itself in the regulation of *MYCN* expression we first compared their expression dynamics. In proliferating A375 cells, *MYCN* expression starts to increase after 72h of culture (Fig 3E), well after that of *LADON*. This delay suggested the regulation of *MYCN* by *LADON* might be indirect. However, the down-regulation of *MYCN* expression after the KD of *LADON* (Fig 3F), and its up-regulation in A375 and A375*Δ*E2 cells transfected with a *LADON*-expressing plasmid (Fig 3G) demonstrated that the *LADON* transcript is indeed promoting *MYCN* expression. The forced expression of *LADON* in A375*Δ*E2 cells thus rescues a molecular defect that is central to their phenotype.

Together these results show that *LADON* promotes the MAT and cell motility in the metastatic melanoma cell line A375 via its impact on the expression of known promoters and suppressors of tumor progression, notably on that of the proto-oncogene MYCN.

### *LADON* expression is dependent on WNT/β-CATENIN signalling

As the MAT has been associated with increased stemness (Taddei et al., 2014) we used an RT2 profiler array (Qiagen) designed to track the expression levels of 84 genes relevant to the stemness of human cancer cells to identify pathways possibly involved in the regulation of *LADON* expression. The analysis was performed on mRNAs extracted from A375 and FO1 cells cultured for 24h and 96h. The comparison of the profiles thus obtained revealed up-regulated and down-regulated genes that were common to both cell lines (Fig 4A, S4A). They included *MYCN*, and a KD experiment showed that in FO1 cells, like in A375 cells, its expression is dependent on that of *LADON* (Fig S4B). They also included *DKK1* and *WNT1*, which encode components of the WNT/*β*-CATENIN signalling pathway. As genes for other components of this pathway (*FZD7*, EPCAM) also showed significant variations (Fig 4A, S4A), and as *MYCN* is known to promote its activation in breast cancer cells (Ma et al., 2010), we tested its involvement. A WB analysis of *β*-CATENIN at regular time points over a 96h culture revealed in both cell lines an increase of the active, unphosphorylated, form of *β*-CATENIN within 48h, whereas total *β*-CATENIN levels remained unchanged (Fig 4B). Treatment with their respective CM for 24h was also found to increase *β*-CATENIN activation. Next, we added recombinant WNT3A to cultured A375 and FO1 cells. RT-PCR analysis showed that a 24h treatment with increasing concentrations of WNT3A led to dose-dependent increases in the expression of AXIN2 – a known target of *β*-CATENIN (Jho et al., 2002) – and *LADON* (Fig 4C), suggesting that this increase might be dependent on WNT/*β*-CATENIN signalling in both cell lines. To confirm this, we used the pharmacological inhibitor XAV, which stabilizes the *β*-CATENIN destruction complex member AXIN2. After 48h of treatment we observed a concomitant decrease in the expression levels of *AXIN2* and *LADON* (Fig 4D). In line with these results, the treatment of A375 and FO1 cells with 60ng/ml of WNT3A also resulted in an increase of MYCN expression (Fig 4E). However, this effect was lost in A375*Δ*E2 cells, in spite of the activation of the pathway in these cells (Fig S4C), demonstrating that the integrity of *LADON*’s locus is required to mediate the entirety of WNT signalling influence on the expression of *MYCN* in the A375 cell line.

**Figure 4.**
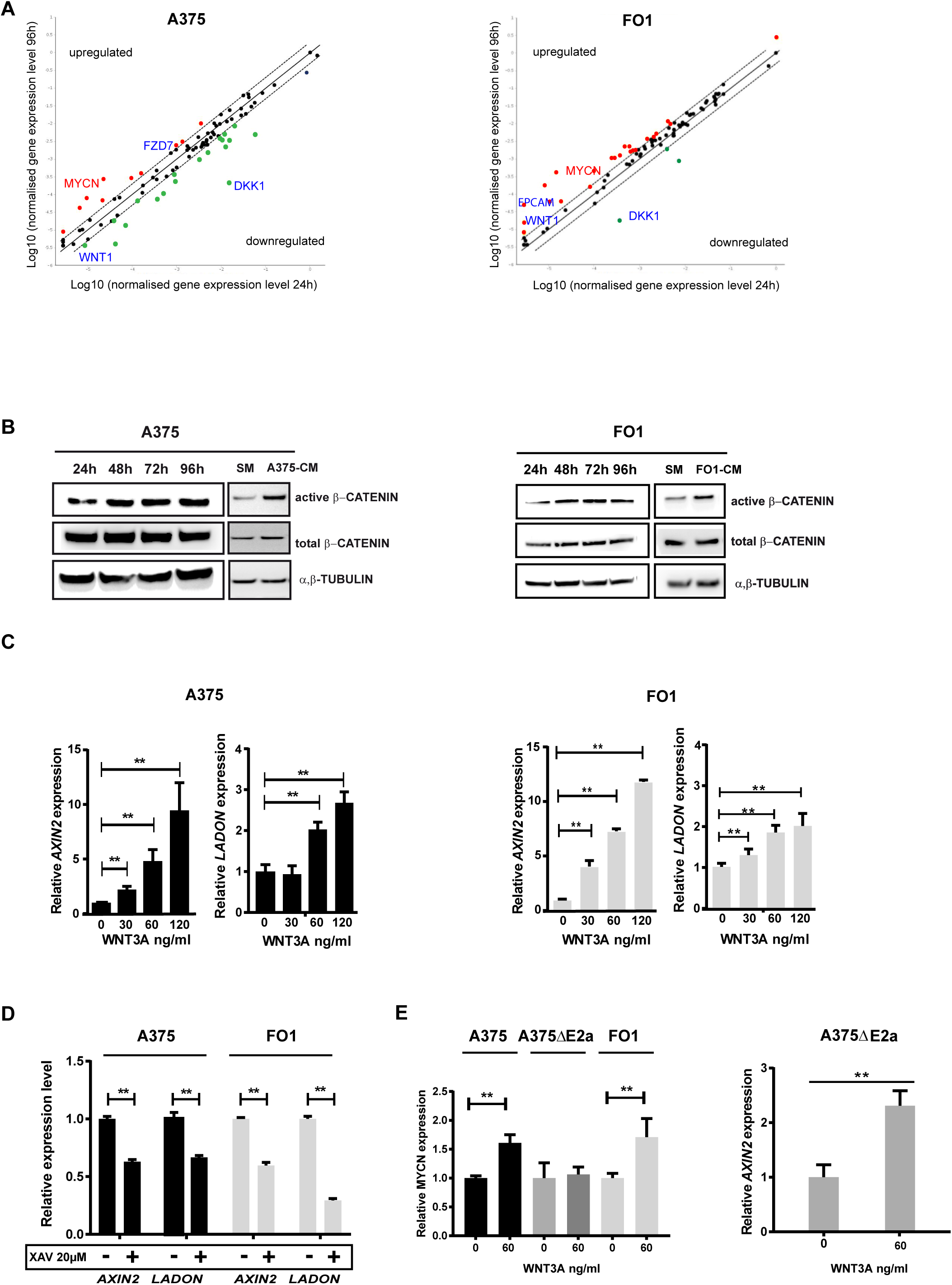
*LADON* expression is dependent on WNT/β-CATENIN signaling. A Scatterplots of the microarray analysis showing how the expression levels of 84 informative genetic markers change in A375 and F01 cells cultured for 96h vs 24h. Upregulated genes are in red, downregulated genes are in green; black dots represent genes that are not differentially expressed. Only changes in expression levels with a fold change > 2 were taken into consideration (see also Table S4A). B Representative immunoblots for active β-CATENIN, total β-CATENIN and α,β-TUBULIN in A375 and FO1 cells cultured in standard medium (SM) for up to 96h, or cultured in SM, A375- conditioned medium (A3751-CM) or FO1-conditioned medium (FO1-CM), for 24h. C RT-qPCR analysis of *AXIN2* and *LADON* expression in A375 or FO1 cells treated with increasing doses of WNT3A. *AXIN2* and *LADON* expression levels were normalized to that of endogenous *RPL13*. Values were then normalized to those obtained for control conditions (0 ng/ml WNT3A), which were set to 1. D RT-qPCR analysis of *AXIN2* and *LADON* expression in A375 or FO1 cells treated with the pharmacological inhibitor XAV939 for 24h. *AXIN2* and *LADON* expression levels were normalized to that of endogenous *RPL13*. Values were then normalized to those obtained for control conditions (0 ng/ml XAV at 24h), which were set to 1. E Left panel: RT-qPCR analysis of *MYCN* expression in A375, A375ΔE2a and FO1 cells treated with 0 (control) or 60 ng/ml WNT3A. Values were normalized to those obtained for controls, which were set to 1. Right panel: RT-qPCR analysis of AXIN2 expression in A375 E2a cells treated with 0 or 60 ng/ml WNT3A, showing that the treatment does activate WNT/ -CATENIN signaling in these cells. The value obtained with WNT3A was normalized to that obtained without, which was set to 1. Histograms display mean values ± SD from a minimum of three independent replicates. P-values were calculated by Student’s t test, * < 0,05, ** < 0.01.

Taken together, these results show that *LADON* expression is positively regulated by WNT/*β*- CATENIN signalling in both the A375 and the FO1 melanoma cell lines and position *LADON* as a key intermediary between WNT/*β*-CATENIN signalling and the activation of *MYCN*.

## Discussion

Our investigations found no evidence that NODAL could influence the behaviour of metastatic melanoma cells *in vitro*, but revealed instead their reliance on the expression of *LADON*, a natural *NODAL* antisense transcript. Our results show that *LADON* is involved in the mechanisms underlying how cells produce pro-metastatic signals and how they respond to them. Experiments conducted in two distinct melanoma cell lines position *LADON* downstream of WNT/*β*-CATENIN signalling and upstream of factors, such as MYCN, known for their implication in the control of tumor progression (Fig 5). Although *LADON*’s mode of action has to be ascertained, our findings support a critical role for its transcript in the network of interactions that governs metastasis in melanoma.

**Figure 5.**
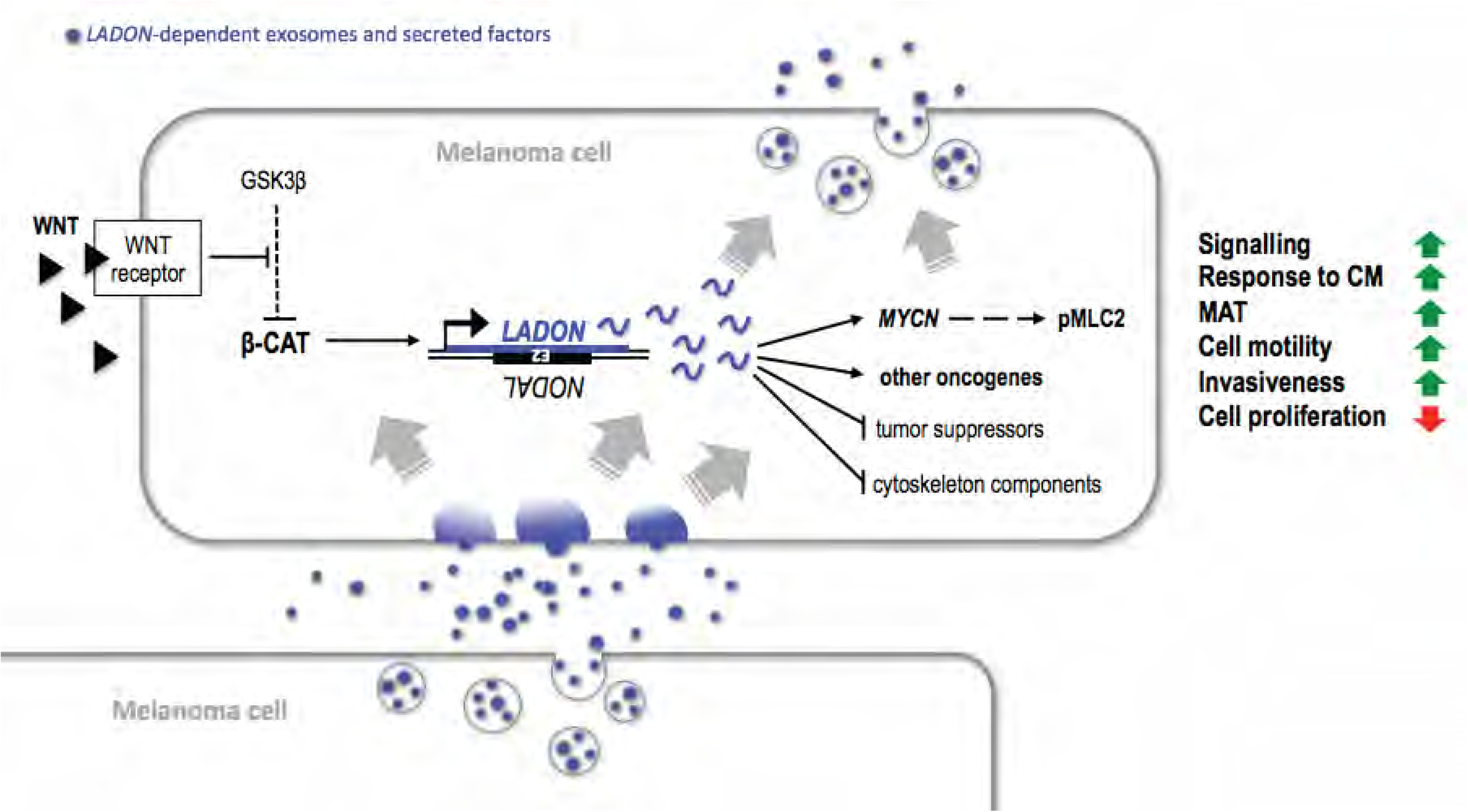
A role for *LADON* in melanoma metastasis. In metastatic melanoma cells, the *LADON* transcript (in blue) promotes the transition from a proliferative cell identity to a less proliferative and more invasive cell identity. In the course of a 4-day culture, a WNT/β-CATENIN-dependent increase of *LADON* expression promotes the expression of known oncogenes and represses that of tumor or metastasis suppressor, which in turn affect the expression of downstream targets, such as cytoskeleton components. *LADON* notably promotes the expression of the proto-oncogene MYCN, which leads to the accumulation of pMLC2, a cytoskeleton component of migratory cells. *LADON* expression also affects the potency of the conditioned medium produced by the cells, presumably via its impact on the exosomes (quantity, content) and the secreted factors they release. *LADON*- dependent exosomes and secreted factors (in blue) promote the progression of exposed melanoma cells towards their more invasive identity. The *LADON* transcript is itself enriched in exosomes.

The absence of *NODAL* transcripts in melanoma cell lines reported here is consistent with the results of other studies (Donovan et al., 2017; Strizzi et al., 2012), as well as with a recent characterization of transcript diversity at the human *NODAL* locus (Findlay & Postovit, 2018). This latter report and our own study show that most exon2-containing RNA species present in melanoma cells are in fact transcribed from the strand opposite to *NODAL*, and correspond to *LADON*. This finding explains some of the inconsistencies found in previous studies of the role of *NODAL* in melanoma, as an exon2-based assay had been used to track the expression of the gene in several of them (Findlay & Postovit, 2018; Hardy et al., 2010; Postovit et al., 2008; Strizzi et al., 2012). The error of mistaking *LADON* expression for that of *NODAL* was compounded by the use of commercial antibodies cross-reacting with non-specific proteins that were erroneously identified as NODAL, as was later demonstrated (Donovan et al., 2017)and our own unpublished results). Donovan et al. make the claim that while there is evidence that TGF*β* and ACTIVIN-A, both signalling via SMAD2,3, do promote tumor progression in melanoma in various ways, no such case can be made for NODAL.

The absence of *NODAL* expression in melanoma cells means that the changes in behaviour we characterized in A375 cells deleted for *NODAL* exon2, the failure to fill the gap in the scratch-wound assay, and the loss in invasiveness, result from the impact of the deletion on the expression of genes other than *NODAL*. The only disturbance we could identify in the expression of the various transcription units in the vicinity of exon2 was that of *LADON*, which expressed a truncated transcript that could no longer respond to inductive treatment. The knockdown of *LADON* with specific siRNAs in both A375 and FO1 cells resulted in a similar loss of invasiveness as the exon2 deletion, even though it was associated with an increase in cell proliferation. This suggests that the failure of exon2-deleted A375 cells to close the gap resulted from a loss of motility so drastic that it was not compensated by their gain in cell number. Finally, the forced expression of *LADON* in exon2-deleted cells reversed the impact of the deletion on the expression of *MYCN*, a key agent of the transition to a more invasive behaviour, thus demonstrating its dependency on the *LADON* transcript itself. These results show that *LADON* is implicated in the transition from a proliferative cell identity to a less proliferative but more invasive cell identity, a change of phenotype inherent to the metastatic process in melanoma (Carreira et al., 2006; Hoek et al., 2006, 2008).

Our finding that the increase in *LADON* expression, which promotes these changes, is dependent on WNT/*β*-CATENIN signalling in at least two melanoma cell lines is at first intriguing. In melanoma, there is evidence of this pathway either promoting (Damsky et al., 2011; Eichhoff et al., 2011a; Gallagher et al., 2013; Murakami et al., 2001; Sinnberg et al., 2011) or suppressing (Arozarena et al., 2011; Bachmann, 2005; Biechele et al., 2012; Chien et al., 2009; Kageshita et al., 2001; Maelandsmo et al., 2003) tumor progression. Most of these discrepancies were explained by context-dependent differences in the expression of downstream effectors of the pathway ( Eichhoff et al., 2011b; Widlund et al., 2002). But two studies have reached opposite conclusions regarding the role of WNT/*β*-CATENIN signalling in melanoma despite being based like ours on the A375 cell line (Biechele et al., 2012; Grossmann et al., 2013) . However, the results underlying these conclusions, showing that *β*-CATENIN signalling promotes a reduction in melanoma cell proliferation (Biechele et al., 2012), but also that it stimulates tumor cell invasion (Grossmann et al., 2013), are in fact not incompatible and are consistent with ours, and lead us to conclude like this latter study that canonical WNT signalling may promote melanoma metastasis.

Another A375-based study recently demonstrated that non-canonical WNT signalling activates Rho-ROCK1/2, and thus promotes melanoma amoeboid invasion, via a FZD7-DAAM1-dependent pathway (Rodriguez-Hernandez et al., 2020). Among its findings it shows that one of the effects of ROCK inhibition is the down-regulation of *CTNNB1* (*β*-CATENIN) and *LADON* (identified as *NODAL* in their qPCR array analysis). This suggests that the surge of ROCK activity that takes place downstream of both the FZD7-DAAM1 pathway and the LADON/MYCN/NDRG1 cascade also promotes *LADON* expression and thus its own persistence. In addition to its negative impact on ROCK1/pMLC2-dependent cell migration NDRG1 is also known to inhibit WNT signalling (Liu et al., 2012; Sun, Zhang, Zheng, et al., 2013), and its repression in A375 cells may thus contribute to the increase in *β*-CATENIN signalling. It seems therefore that in A375 cells at least, canonical WNT signalling promotes invasion and its own activity, and even appears to synergize with non-canonical WNT signalling to do so.

Whether it is suppressing or promoting its progression, the importance of WNT/*β*-CATENIN signalling in melanoma is not in doubt. As a target of the pathway and a regulator of downstream oncogenes and tumor suppressors, *LADON* could provide a valuable point of entry for approaches aimed at derailing metastatic processes. It will be important in future studies to find out whether *LADON* is also a target of WNT/*β*-CATENIN signalling in other cancers where the pathway is known to play an important role in tumor progression.

There are now multiple examples of the implication of lncRNAs in tissue physiology and in the misregulation of cellular processes, which often result in cancer (Huarte, 2015; Iyer et al., 2015; Schmitt & Chang, 2016) . This implication is the reason for the interest in lncRNAs as biomarkers for diagnostic and prognosis, but also as potential targets for therapy or as tools to correct defective gene expression underlying cancer and other pathologies (Slaby et al., 2017; Wahlestedt, 2013). How *LADON* is regulating the expression of its downstream effectors is as yet unclear. It takes 24h to detect a significant change in the expression of *MYCN* after that of *LADON* has been altered, a delay implying that this regulation may not involve a direct interaction between *LADON* and this effector. Even at its peak the expression of *LADON* never reaches a level that would allow it to be an effective miRNA sponge (Denzler et al., 2014, 2016). An interesting feature of the transcript is the presence in its 3’region of the short interspersed elements (SINEs) Alu and MIR. The presence of Alu elements in lncRNAs has been associated with the capacity of such transcripts to regulate RNA transcription, decay or splicing (Gong & Maquat, 2011; Holdt et al., 2013; Hu et al., 2016), while the presence of other SINEs has been associated with the regulation of other processes, such as translation (Carrieri et al., 2012; Johnson & Guigó, 2014). Interestingly, it has been shown in neuroblastoma cells that *β*-CATENIN signalling can modulate *MYCN* expression via its impact on the stability of the *MYCN* mRNA (Duffy et al., 2014). Further investigations will be necessary to assess whether it is the presence of SINEs in the *LADON* transcript that conditions its capacity to modulate the expression of *MYCN* expression, as well as that of other genes, in melanoma cells.

To conclude, we identified a specific role for *LADON* in the regulatory cascade that via MYCN and other factors allows melanoma cells to become more invasive. However, the evidence gathered so far, notably its impact on the expression of oncogenes and tumor suppressors known to be associated with processes other than cell motility and invasiveness, suggests its implication in tumor progression is likely to be broader. *LADON* expression has so far only been reported in melanoma and breast cancer cell lines (Findlay & Postovit, 2018), and this study), but is likely to be present in other cancer cell lines. There is evidence that it is present in healthy human cell types and tissues, where we currently have no indication of its function. Most of these cells, whether cancerous or not, do not express *NODAL*, but for those that do, the possibility remains that *LADON* then has the ability to modulate NODAL signalling. Future studies of *LADON* should therefore focus on its function and mode of action in a variety of contexts.

## Material and Methods

### Key Resource Table

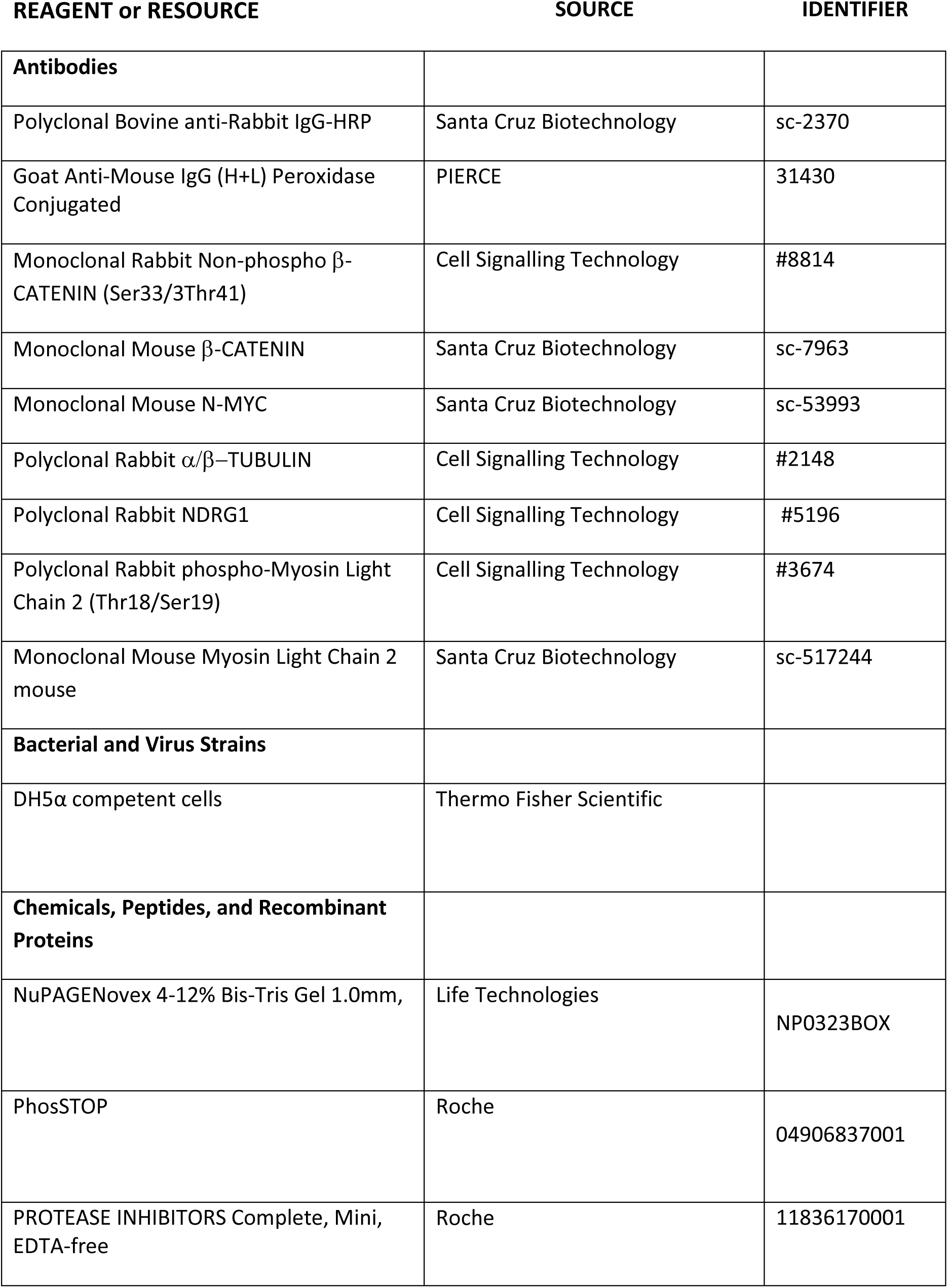
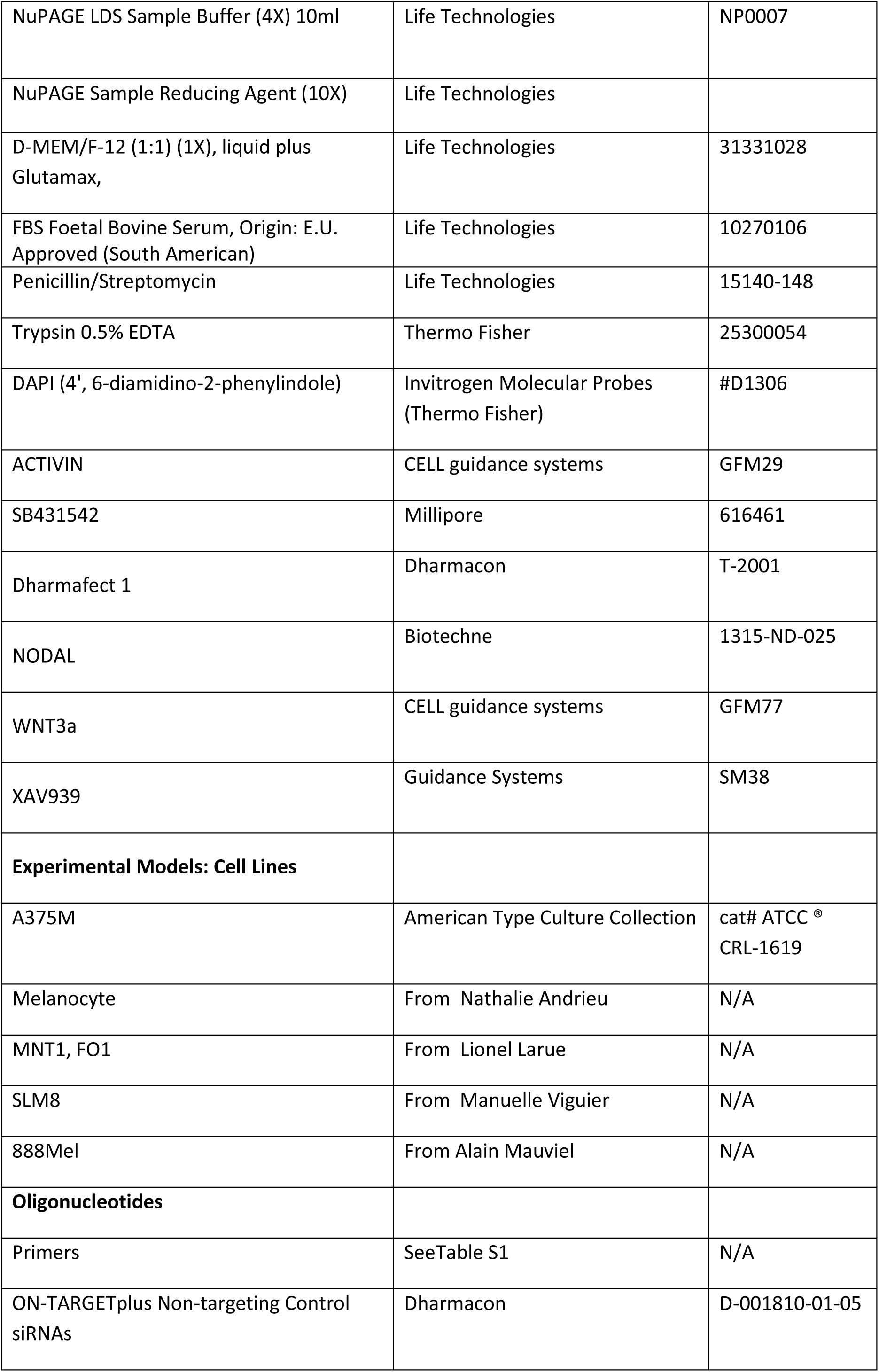
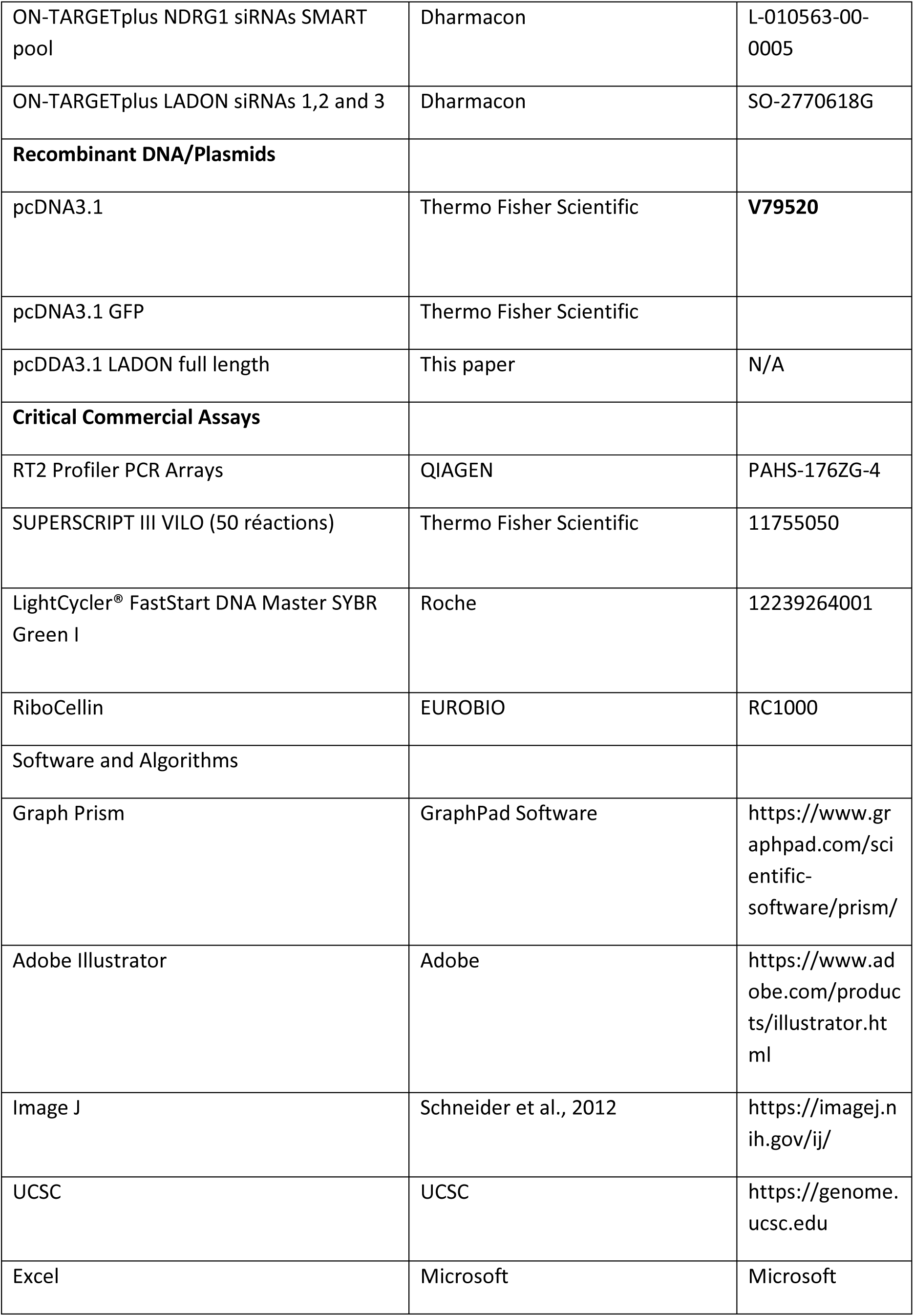

### Plasmids

pcDNA3.1 and GFP pcDNA 3.1 are from Thermo Fisher scientific. LADON corresponding to AK001176.1, 1725bp was synthetized and inserted in pCDN3.1 (Genscript).

### Cell lines and cell culture

The melanoma cell line A375M was purchased from ATCC. The normal melanocyte cell line was kindly provided by Nathalie Andrieu (Centre de Recherches en Cancérologie de Toulouse). The SLM8 cell line, kindly provided by M Viguier, is derived from a lymph node metastasis. The MNT1 and FO1 melanoma cell lines were a gift from Lionel Larue (Institute Curie CNRS UMR3347, INSERM U1021, Institute Curie). The 888Mel cell line, a gift from A. Mauviel (Institut Curie/CNRS UMR 3347/INSERM U1021), is derived from the lung metastatic WM793 melanoma cell line. Cells were grown in DMEM/F12 Glutamax (Invitrogen, Cergy-Pontoise, France) supplemented with antibiotics and 10% fetal calf serum (FCS) in a 5% CO2 atmosphere.

To prepare conditioned medium, melanoma cell lines were seeded at 70% confluence and then incubated for 72h in DMEM medium with or without FCS, as stated. This melanoma-conditioned medium was collected, centrifuged at 11000g for 15 min to remove cell debris and stored at -80°C.

The source and identifier of all the plasmids and cell lines are listed in the Key Resources Table.

### Transmigration assay

Melanoma cells (1×10^5^) were seeded on the upper compartments of 2 mg ⁄ ml type I collagen-coated culture inserts (8 µm pores -Greiner Bio-One SAS, Courtaboeuf, France). DMEM supplemented with 10% FCS was used as a chemoattractant. A375 and A375*Δ*E2 cells were allowed to migrate at 37°C and 5% CO2 for 24h00. Non-migrating cells on the upper face of the filter were removed by gently scraping them off using a cotton swab. Cells on the lower face were washed in PBS, fixed with 4% formaldehyde for 10 min and washed in PBS. Nuclei were then labeled with Hoechst for 5 min and washed again. Migrating cells were observed under an epifluorescence microscope using a 10x magnification. Five to ten pictures of adjacent fields of the central zone of each Transwell were taken. Fluorescence was quantified with the IMAGEJ software (US National Institutes of Health, Bethesda, MD, USA). Histograms display the data obtained from three independent experiments, and all the experiments were performed in duplicate. P-values were calculated by means of an ANOVA. Data are displayed as normalized results, where the nuclei of melanoma cells are set to 1 (or 100%) when grown in control condition.

### A375 staining

For immunofluorescence experiments, A375 cells were seeded onto glass coverslips in twelve-well plates and incubated overnight. Next, cells were fixed in 4% formaldehyde solution and incubated with Alexa Fluor 647-coupled phalloidin to visualize F-actin, and Hoechst 33342 to label nuclei. Cover slips were then further washed in PBS, treated with Fluoromount-G anti-fade (Southern Biotech), and analyzed by confocal microscopy. The shape of A375 cells (n>20), elongated or round, was analyzed by IMAGEJ software, and the proportion of each corresponding cell type was quantified.

### Wound healing/scratch assay

Cell migration was examined using a wound-healing assay. In brief, 0.2×10^6^ cells were seeded in a well of twelve-well plates and at confluence a scratch wound was made with a 10 µl pipette tip, and then washed twice with PBS to remove cell debris. Wells were photographed under phase-contrast microscopy (time=0) while cells were allowed to migrate into the scratch wound area for up to 18h at 37°C 5% C02 atmosphere using an Essenbio IncuCyte apparatus. Speed of closure area was calculated overtime by IMAGEJ software. Data are represented as a ratio of migratory cells normalized to A375 cells migration in control condition. All experiments were performed in duplicate.

### Western blot

Briefly, cells were lysed using RIPA buffer (50 mMTris–HCl, 10 mM MgCl_2_, 20% Glycerol and 1% Triton X-100) containing protease and phosphatase inhibitors (Roche). Protein concentration was determined by BCA assay (Pierce) and 10-30 μg used for Western blot analysis. Western blot samples were combined with NuPAGE LDS sample buffer (Life Technologies) containing 0.05% β-mercaptoethanol, incubated at 95°C for 5 minutes and then resolved on SDS-PAGE (12% acrylamide). The resolved proteins were blotted onto nitrocellulose membrane and blocked with 5% milk or BSA (weight/volume). These were then incubated with primary antibodies overnight at 4°C, followed by an incubation at room temperature with the respective horseradish peroxidase-conjugated secondary antibodies for 30 minutes. Details of the antibodies used are shown in the ‘antibodies and reagents’ section. Immunoreactive proteins were detected by enhanced chemiluminescence using a ChemiDoc XRS Molecular Imager (BioRad). Blots were quantified using the software available with this instrument. For all figures, representative blots are shown from replicate experiments.

### CRISPR/CAS9 Genome Editing

CRISPR/Cas9 genome editing was performed with GeneArt CRISPR Nuclease Vector Kit according to manufacturer’s instructions (Life Technology). To select the target sequence for genome editing, the genomic sequences surrounding *NODAL* exon2 were submitted to an online CRISPR Design Tool (http://crispr.mit.edu/). Two target sites were selected upstream and downstream of this sequence. The oligonucleotides used to construct gRNAs for the human *NODAL* gene exon2 (deletion of 698 pb) are listed in Table S1.

The ds oligonucleotides generated were cloned into the GeneArt CRISPR Nuclease Vector. Competent E. coli cells were transfected with 3 μL of ligation reaction, and then 50 µL from the transformation reaction was spread on a pre-warmed LB agar plate containing 100 µg/mL ampicillin. Plates were incubated overnight at 37°C. The identity of the ds oligonucleotide insert in positive transformants was confirmed by sequencing. For A375 cell line transfection, the cationic lipid-based Lipofectamine 2000 Reagent was used. A375 cells positive for the transfection were sorted by FACS using OFP, a fluorescent protein present in the GeneArt CRISPR Nuclease Vector. Genotyping was performed on 96 clones to detect the deletion. Primers used to assess the efficacy of the CRISPR/Cas9 deletion are listed in Table S1.

Length of non-deleted amplicon: 1141 pb Length of deleted amplicon: 445 pb

### siRNAs

siRNAs against NDRG1 (mix of 4), LADON (two independents) and negative-control RNA were chemically synthesized (Dharmacon Research, Lafayette, USA). Synthetic siRNAs were transfected with Ribocellin Transfection Reagent (Eurobio) according to the manufacturer’s instructions.

### RNA extraction, reverse transcription (RT) and Quantitative PCR (Q-PCR)

For PCR analysis, 10^5^ transfected GFP-positive cells were sorted by FACS analysis and collected into RNAse-free tubes. Total RNA extraction, cDNA synthesis and Q-PCR were performed as described previously (Legent et al., 2006). For each gene and for a given RT PCR, values were normalized to the level of expression of the reference genes *RLP13* and *GAPDH*. No significant differences in the final ratio were found between the two reference genes; therefore, only *RPL13* was used for normalization. Primers and annealing temperatures for all genes are indicated in Table S1. For each gene, the values were averaged over at least three independent measurements. Three independent RNA isolations were performed for all experiments.

Sequences for all siRNAs and oligos used in this study can be found in Table S1.

### RT^2^ Profiler PCR Array-

The Human Cancer Stem Cells RT² Profiler PCR Array from Qiagen was used to profile the expression of 84 genes linked to cancer stem cells (CSCs) in the A375 and FO1 cell lines after 24h and 96h of culture, according to the manufacturer’s instructions.

### LC-MS/MS acquisition

For proteomics analysis two A375*Δ*E2 clones (a and d) and two parental clones (A375a and b) were used.

Protein extracts (60µg) were precipitated with acetone at -20°C; the protein extracts were then incubated overnight at 37°C with 20 μl of 25mM NH_4_HCO_3_ containing sequencing-grade trypsin (12.5μg/ml, Promega). The resulting peptides were desalted using ZipTip µ-C18 Pipette Tips (Millipore) and analyzed either in technical triplicates or individually by a Q-Exactive Plus coupled to a Nano-LC Proxeon 1000 equipped with an easy spray ion source (all from Thermo Scientific). Peptides were separated by chromatography with the following parameters: Acclaim PepMap100 C18 pre-column (2cm, 75μm i.d., 3μm, 100Å), Pepmap-RSLC Proxeon C18 column (50cm, 75μm i.d., 2μm, 100 Å), 300 nl/min flow rate, gradient from 95 % solvent A (water, 0.1% formic acid) to 35% solvent B (100% acetonitrile, 0.1% formic acid) over a period of 98 min, followed by a column regeneration for 23 min, giving a total run time of 2 hrs. Peptides were analyzed in the Orbitrap cell, in full ion scan mode, at a resolution of 70,000 (at *m/z* 200), with a mass range of *m/z*375-1500 and an AGC target of 3×10^6^. Fragments were obtained by high collision-induced dissociation (HCD) activation with a collisional energy of 30%, and a quadrupole isolation window of 1.4 Da. MS/MS data were acquired in the Orbitrap cell in a Top20 mode, at a resolution of 17,500 with an AGC target of 2×10^5^, with a dynamic exclusion of 30 sec. MS/MS of most intense precursor were firstly acquired. Monocharged peptides and peptides with unassigned charge states were excluded from the MS/MS acquisition. The maximum ion accumulation times were set to 50 ms for MS acquisition and 45 ms for MS/MS acquisition.

### LC-MS/MS data processing

The LC-MS/MS.raw files were processed using the Mascot search engine (version 2.5.1) coupled to Proteome Discoverer 2.2 (Thermo Fisher Scientific) for peptide identification with both a custom database and the database Swissprot (from 2017) with the *Homo sapiens* taxonomy. The following post-translational modifications were searched on proteome Discoverer 2.2: Oxidation of methionine, acetylation of protein N-term, phosphorylation of serine/threonine and phosphorylation of tyrosine. Peptide Identifications were validated using a 1% FDR (False Discovery Rate) threshold calculated with the Percolator algorithm.

### Data collection and analysis

BigWig coverage files for the plus and minus strands were retrieved for the keratinocytes, and melanocytes in the skin02 sample (ENCODE project). The data retrieved from the National Center for Biotechnology Information Gene Expression Omnibus were accessible through GEO series number GSE78652 for A375cells, GSE16256 for H1 cells, for the A375 melanoma line, H1 cells, the bam files for the samples were first merged before deriving the bigWig coverage file.

The HG-U133 plus 2 arrays (Affymetrix) data presented are accessible through GEO Series accession GSE35388 analyzed by GEO2R at the National Center for Biotechnology Information Gene.

### Statistical analysis

Data was collected and graphed using GraphPad Prism software. A 1-way ANOVA or two-sided student’s t-test was used to determine statistical significance where appropriate. P-values of less than 0.05 were considered statistically significant. Information about the statistics used for each experiment, including sample size, experimental method, and specific statistic test employed, can be found in the relevant figures or figure legends.

## Author Contributions

AD, JC, DF conceived the research. JC and DF obtained funding. JC and DF oversaw the experiments and data analysis. AD and DF carried out the experiments. AD analyzed the data. SD contributed to transmigration assays and RT-qPCR analysis. SC contributed to transmigration assays and RT-qPCR analysis. CB contributed to PCR, RT-qPCR analysis and CRISPR cloning. SH contributed to WB analysis and manuscript editing. FD contributed to the transmigration assays and provided essential reagents and cell lines. JC, AD and DF wrote the manuscript.

## Acknowledgements

We thank Nathalie Andrieu for the melanocyte cell line, Manuelle Viguier for the SLM8 cell line, Lionel Larue for the MNT1 cell line, Alain Mauviel for the 888Mel cell line and Delphine Delacour for antibodies. We thank Daniel Constam, Lionel Larue, Claire Rougeulle, Delphine Delacour and Vanessa Ribes for discussion and suggestions. We thank the Proteomics Core Facility at Institut Jacques Monod, notably T. Léger and C. Garcia, for the LC-MS/MS experiments, and the Région Ile-de-France (SESAME 2013 Q-Prot-B&M -LS093471), the Paris-Diderot University (ARS 2014-2018), and CNRS (Moyens d’Equipement Exceptionnel INSB 2015) for funding part of the LC-MS/MS equipment.

We also acknowledge the ImagoSeine Core Facility of Institut Jacques Monod, member of IBISA and of the France-Bioimaging (ANR-10-INBS-04) infrastructures.

Work in the Collignon lab was supported by grants from INCa (2014-1-PL BIO-01) and GEFLUC (2015-2017), and by Centre National de la Recherche Scientifique (CNRS).

Chiara BRESESTI was supported by the EUR G.E.N.E. (#ANR-17-EURE-0013), which is part of the Université de Paris IdEx (#ANR-18-IDEX-0001) funded by the French Government.

