## Supplementary material for "*LADON*, a natural antisense transcript of *NODAL*, promotes an invasive behaviour in melanoma cells": Fig. 1S

Figure S1.

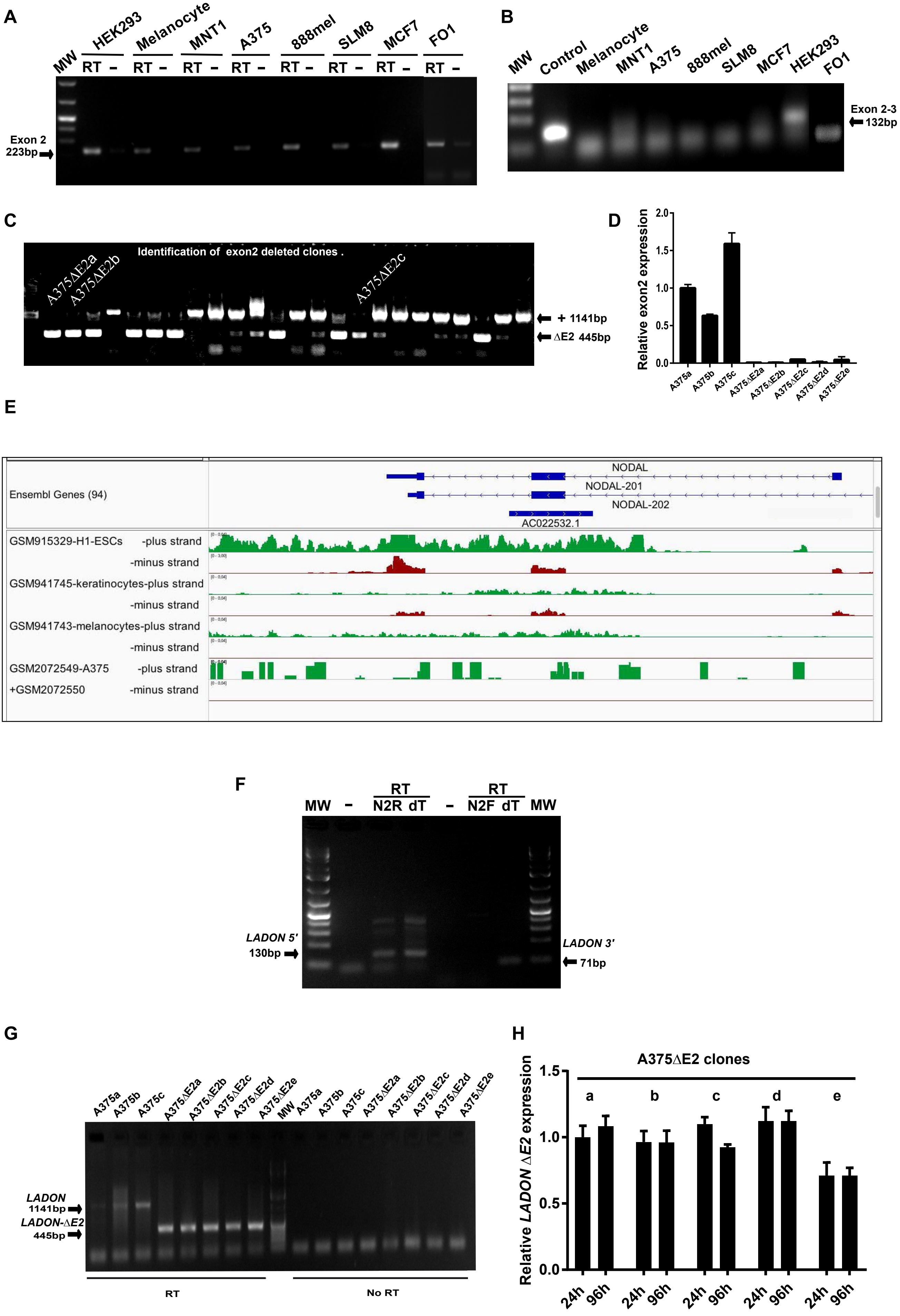

A
MW
HEK293
Melanocyte
MNT1
A375
888mel
SLM8
MCF7
FO1
RT –
Exon 2
223bp
B
MW
Control
Melanocyte
MNT1
A375
888mel
SLM8
MCF7
HEK293
FO1
Exon 2-3
132bp
C
Identification of exon2 deleted clones .
A375ΔE2a
A375ΔE2b
A375ΔE2c
+ 1141bp
ΔE2 445bp
D
Relative exon2 expression
2.0
1.5
1.0
0.5
0.0
A375a
A375b
A375c
A375ΔE2a
A375ΔE2b
A375ΔE2c
A375ΔE2d
A375ΔE2e
E
Ensembl Genes (94)
NODAL
NODAL-201
NODAL-202
AC022532.1
GSM915329-H1-ESCs -plus strand
-minus strand
GSM941745-keratinocytes-plus strand
-minus strand
GSM941743-melanocytes-plus strand
-minus strand
GSM2072549-A375 -plus strand
+GSM2072550 -minus strand
F
MW
–
RT
N2R dT
–
RT
N2F dT
MW
LADON 5'
130bp
LADON 3'
71bp
G
A375a
A375b
A375c
A375ΔE2a
A375ΔE2b
A375ΔE2c
A375ΔE2d
A375ΔE2e
MW
LADON
1141bp
LADON-ΔE2
445bp
RT
No RT
H
A375ΔE2 clones
a
b
c
d
e
Relative LADON ΔE2 expression
1.5
1.0
0.5
0.0
24h
96h

**Figure S1. *LADON* is a *NODAL* natural antisense transcript expressed in a variety of cell lines**

A RT-PCR with the primers N2R and N2F to detect the transcription of exon2 in a panel of cell lines: embryonic kidney (HEK293), melanocytes, non-metastatic melanoma (MNT1), metastatic melanoma (A375, 888mel, SLM8), breast cancer (MCF7). The expected 223bp band is detected in all reverse-transcribed samples (noted RT), but not in samples without reverse transcriptase (noted -).

B RT-PCR with primers spanning the *NODAL* exon 2-3 boundary in the same cell lines detected the expected 132bp band only in the HEK293 cell line and in the positive control, which is a plasmid containing a cDNA of *NODAL*.

C PCR amplification of the genomic region targeted by the CRIPSR/Cas9-mediated deletion of *NODAL* exon2 in A375 cells. In the 24 independent clones analyzed, amplification of the intact locus yields a 1141nt-long band, while that of the deleted one (ΔE2) yields a 445nt-long band.

D RT-qPCR of *NODAL* exon2 with the N2F/N2R primer pair detects the presence of a transcript in 3 independent A375 parental clones (A375a-c), but not in the 5 independent mutant clones A375ΔE2a to e.

E Mapping of strand-specific expression data at the human *NODAL* locus in embryonic stem cells (ESC), in cell lines representative of skin cell types (keratinocytes, melanocytes), and in the A375 melanoma cell line. Read coverage is displayed in red for the minus strand (from which *NODAL* is transcribed) and in green for the plus strand (from which *LADON* is transcribed). The exonic structure of the two genes is shown.

F PCR amplification of reverse-transcribed total A375 RNA yields bands for *LADON* from cDNAs primed with an oligo dT or the reverse primer N2R, but not from cDNAs primed with the forward primer N2F. No band is amplified in samples without reverse transcriptase (noted -).

G RT-PCR using primers spanning the entire exon2 detects the presence of a transcript, corresponding to *LADON-ΔE2*, a truncated version of *LADON*, in the exon2-deleted clones A375ΔE2a to e.

H RT-qPCR analysis of *LADON-ΔE2* expression in five independent A375ΔE2 clones detects no increase over the course of the culture.
