## Supplementary material for "*LADON*, a natural antisense transcript of *NODAL*, promotes an invasive behaviour in melanoma cells": Fig. 2S

### Figure S2

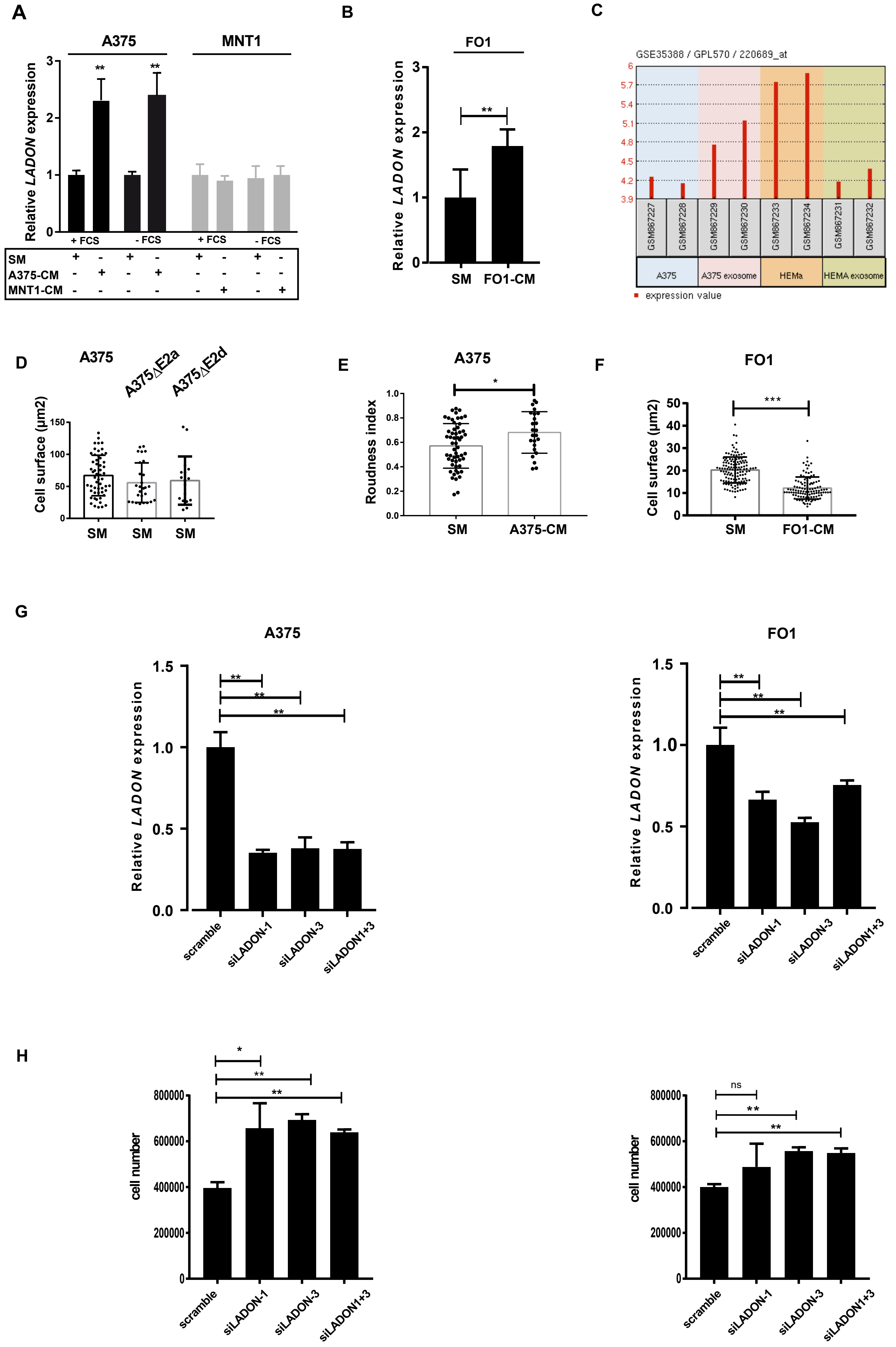
A
A375
MNT1
Relative LADON expression
+ FCS
- FCS
SM
A375-CM
MNT1-CM
B
FO1
Relative LADON expression
SM
FO1-CM
C
GSE35388 / GPL570 / 220689_at
GSM867227
GSM867228
GSM867229
GSM867230
GSM867233
GSM867234
GSM867231
GSM867232
A375
A375 exosome
HEMa
HEMA exosome
expression value
D
A375
A375△E2a
A375△E2d
Cell surface (µm2)
SM
E
A375
Roudness index
SM
A375-CM
F
FO1
Cell surface (µm2)
SM
FO1-CM
G
A375
FO1
Relative LADON expression
scramble
siLADON-1
siLADON-3
siLADON1+3
H
cell number
ns

**Figure S2. Causes and consequences of changes in *LADON* expression**

A RT-qPCR analysis of *LADON* expression in A375 and MNT1 cells cultured for 24h in standard culture medium (SM), in A375-conditioned medium (A375-CM) or in MNT1- conditioned medium (MNT1-CM), with or without fetal calf serum (FCS).

B RT-qPCR analysis of *LADON* expression in FO1 cells cultured for 24h in SM or in FO1-conditoned medium (FO1-CM).

C *LADON* expression levels measured in melanoma cells (A375), in normal melanocyte (HEMa), and in their exosomes ( GSE35388 dataset on the Affymetrix Human Genome U133 Plus 2.0 Array analysis with the 220689_at probe set).

D The histogram shows cell surface measurements of A375, A375ΔE2a and A375ΔE2d cells cultured for 24h in SM.

E The histogram shows roundness index measurements of A375 cells cultured for in SM or A375-SM.

F The histogram shows cell surface measurements of FO1 cells cultured for 24h in SM or FO1-CM.

G RT-qPCR analysis of *LADON* expression in A375 or FO1 cells treated with the siRNAs scrambled, siLADON-1, siLADON-3, or siLADON-1 and -3. Values are normalized to that obtained with the scrambled siRNA, which is set to 1.

H Comparison of cell counts in A375 or FO1 cells cultured for 72h in the presence of siRNAs scrambled, siLADON-1, siLADON-3, or siLADON-1 and -3.

Histograms display mean values ± SD from a minimum of three independent replicates. P-values were calculated by Student's t test, * < 0.05, ** < 0.01.
