## Supplementary material for "*LADON*, a natural antisense transcript of *NODAL*, promotes an invasive behaviour in melanoma cells": Fig. 4S

A

### Macro array analysis 24h vs 96h

### A375

| Gene Symbol | Fold Regulation |
|---|---|
| MYCN | 12,25 |
| FOXP1 | 8,57 |
| NANOG | 6,5 |
| DLL1 | 3,26 |
| DACH1 | 3,25 |
| MUC1 | 3,06 |
| IL8 | 2,86 |
| ATXN1 | 2,55 |
| TWIST1 | 2,53 |
| FZD7 | 2,36 |
| EGF | -2,01 |
| FGFR2 | -2,14 |
| WNT1 | -2,36 |
| PLAUR | -2,42 |
| HPRT1 | -2,43 |
| PLAT | -2,43 |
| CHEK1 | -2,55 |
| SAV1 | -2,96 |
| ACTB | -3,08 |
| JAG1 | -3,29 |
| KLF4 | -3,35 |
| WWC1 | -3,99 |
| YAP1 | -4,68 |
| KITLG | -4,76 |
| ABCG2 | -5,01 |
| NOS2 | -5,21 |
| MYC | -5,88 |
| PROM1 | -10,27 |
| DKK1 | -12,13 |

### FO1

| Gene Symbol | Fold Regulation |
|---|---|
| MUC1 | 28,34 |
| BMP7 | 21,71 |
| EPCAM | 17,33 |
| THY1 | 5,72 |
| KIT | 5,58 |
| ITGA2 | 4,77 |
| MYCN | 4,53 |
| POU5F1 | 4,32 |
| MERTK | 3,97 |
| ATXN1 | 3,77 |
| FOXP1 | 3,43 |
| KLF17 | 3,28 |
| GATA3 | 2,94 |
| EGF | 2,74 |
| WNT1 | 2,71 |
| GAPDH | 2,57 |
| TWIST1 | 2,55 |
| JAK2 | 2,5 |
| ALDH1A1 | 2,45 |
| ATM | 2,21 |
| KLF4 | 2,19 |
| ALCAM | 2,06 |
| ITGA4 | 2,04 |
| FGFR2 | 2,02 |
| AXL | -2,07 |
| IL8 | -8,34 |
| DKK1 | -19,97 |

| |
|---|
| 30 |
| 20 |
| 15 |
| 10 |
| 8 |
| 6 |
| 5 |
| 4 |
| 3 |
| 2 |
| -2 |
| -3 |
| -4 |
| -5 |
| -6 |
| -8 |
| -10 |
| -12 |
| -20 |

Asymmetric division

WNT/β-catenin

B

FO1

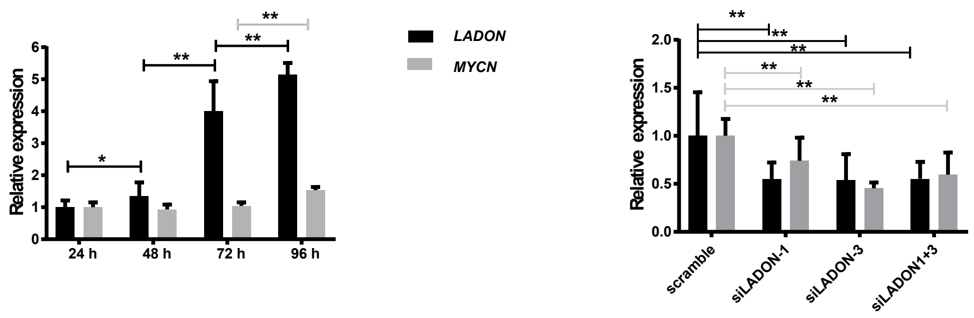

**Figure S4. Gene expression changes associated with culture duration in A375 and FO1 cells**

A A375 and FO1 cells were cultured for 24h and 96h, and differences in the expression of 84 genes relevant to stemness were revealed using an RT2 profiler array. The list of differentially expressed genes and their respective fold change are detailed with red to green color coding. Genes highlighted in pink or yellow are associated with asymmetric division or WNT/B catenin signaling pathways respectively.

B Left panel: RT-qPCR analysis of *LADON* and *MYCN* expression in FO1 cells cultured for 24h, 48h, 72h or 96h in SM ($n \geq 3$). Right panel: RT-qPCR analysis of *LADON* and *MYCN* expression in FO1 cells treated with the siRNAs scrambled, siLADON-1, siLADON-3, or siLADON-1 and -3. Values were normalized to those obtained with the scrambled siRNA, which were set to 1.

Histograms display mean values ± SD from a minimum of three independent replicates. P-values were calculated by Student's t test, * < 0,05, ** < 0.01.
