## Supplementary material for "*LADON*, a natural antisense transcript of *NODAL*, promotes an invasive behaviour in melanoma cells": Table 1S

Table S1.The primers used for qRT-PCR, CRISPR/CAS9 genome editing and siRNA. Related to STAR Methods.

| Amplicon | Sequence (5'-3') | Amplicon size |
|---|---|---|
| CRISPR/CAS9 Exon 2 Deletion 5' | F- ATCCACTGCCACATCTGGGTGTTTT<br>R- ACCCAGATGTGGCAGTGGATCGGTG | |
| CRISPR/CAS9 Exon 2 Deletion3' | F- GACCAACCATGCATACATCCGTTTT<br>R- GGATGTATGCATGGTTGGTCCGGTG | |
| ladon $\Delta E2$detection | F- ATCTTGGAAGGGGGACTGGAA<br>R- TCCTGGTACATTGGAGGTGCTT | 1141- 445 |
| Nodal Exon2 | N2F- CTGCTTAGAGCGGTTTCAGATG<br>N2R- CGAGAGGTTGGAGTAGAGCATAA | 224 |
| Nodal Exon 2-3 | F- AGCAGTACAACGCCTATCGC<br>R- AACAAGTGGAAGGGACTCGG | 132 |
| LADON 5' | L1F- CAAAGCTAGAGCCCTGTCCC<br>L1R- AGGGCGAGTGTCCTAATCCT | 130 |
| LADON 3' | L4F- AAGCAAACGTCCAGTTCTGCC<br>L4R- TCACCCTCCTTCCTTCTTGGT | 71 |
| GAPDH | F- GAAGGTCGGAGTCAACGGATT<br>R- TGACGGTGCCATGGAATTTG | |
| RPL13 | F- GGAAGTACCAGGCAGTGACAGC<br>R- CTTCTCGGCCTGTTTCCGTAG | |
| Actinb | F- GAGCTACGAGCTGCCTGAC<br>R- GCACTGTGTTGGCGTACA | |
| MYCN | F- CACGTCCGCTCAAGAGTGTC<br>R- GTTTCTGCGACGCTCACTGT | |
| Axin-2 | F- GCTGACGGATGATTCCATGT<br>R- ACTGCCCACACGATAAGGAG | |
| NDRG1 | F- CAAGATCTCAGGATGGACC<br>R- GACCACTTCCACGTTACTC | |
| siRNA NODAL-AS1 | 1) CAGCUAAUGAGGAGUCAAAUU<br>2) CAUUGGAGGUGCUUGAGU<br>3) ACAGGAACAUAGUCAUUUAUU | |
| siRNA NDRG1 | 1) GCAGGCGCCUACAUCCUAA<br>2) GAAGAAAUGCAGAGUAACG<br>3) GUGGAGGGCCUUGUCCUUA<br>4) UCGUGAGGUGAAGCCUUU | |
| probe set 220689_at | CTCAGTCAGTGTCACAACCACCATA<br>CAACCACCATAGCTCTTGCTGGAGG<br>CTTGCTGGAGGCCTTACCTTATATA<br>GATTTTCGAGTCCCTTTTAGAGATG<br>AGTAAACCTGGATTTGCCCAACTGC<br>AGCCCAGACTCCGAATTACTGTTCT<br>GTCCCCCTTCCAAGATAACTTGCAC<br>CTTGCACAGCCAGAGGACAGTCAGT<br>AAATGTGATGAGTTGCATCTCCCCC<br>CTCCTCCCATACATGAAGTCTGGCC<br>CCCAGCCACTCGAGAATCATTTGAA | |
